# Contribution of deep soil layers to the transpiration of a temperate deciduous forest: quantification and implications for the modelling of productivity

**DOI:** 10.1101/2022.02.14.480025

**Authors:** Jean Maysonnave, Nicolas Delpierre, Christophe François, Marion Jourdan, Ivan Cornut, Stéphane Bazot, Gaёlle Vincent, Alexandre Morfin, Daniel Berveiller

## Abstract

Climate change is imposing drier atmospheric and edaphic conditions on temperate forests. Here, we investigated how deep soil (down to 300 cm) water extraction contributed to the provision of water in the Fontainebleau-Barbeau temperate oak forest over two years, including the 2018 record drought. Deep water provision was key to sustain canopy transpiration during drought, with layers below 150 cm contributing up to 60% of the transpired water in August 2018, despite their very low density of fine roots. We further showed that soil databases used to parameterize ecosystem models largely underestimated the amount of water extractable from the soil by trees, due to a considerable underestimation of the tree rooting depth. The consensus database established for France gave an estimate of 207 mm for the soil water holding capacity (SWHC) at Fontainebleau-Barbeau, when our estimate based on the analysis of soil water content measurements was 1.9 times as high, reaching 390±17 mm. Running the CASTANEA forest model with the database-derived SWHC yielded a 350 gC m^−2^ y^−1^ average underestimation of annual gross primary productivity under current climate, reaching up to 700 gC m^−2^ y^−1^ under climate change scenario RCP8.5. It is likely that the strong underestimation of SWHC that we show at our site is not a special case, and concerns a large number of forest sites. Thus, we argue for a generalisation of deep soil water content measurements in forests, in order to improve the estimation of SWHC and the simulation of the forest carbon cycle in the current context of climate change.

**Highlights:**

- Forest-atmosphere carbon exchanges remained insensitive to record drought.
- Deep soil (150-300 cm) provisioned up to 60% of the water transpired by the forest during drought.
- Soil databases were underestimating soil water holding capacity by a factor of two.
- Simulated forest productivity is strongly sensitive to soil water holding capacity parameter.
- Deep soil water content measurements are urgently needed to correctly estimate the soil water holding capacity.

## Introduction

Evaporating water is vital for trees. The process of evapotranspiration plays a central role in the thermoregulation of leaves, the acquisition and transport of nutrients from the soil to plant organs and is an inescapable consequence of the stomatal opening for the acquisition of carbon from the atmosphere. In plant communities, orders of magnitude for the amount of water transpired per unit carbon gained through photosynthesis range from 190 grams of water per unit gram of carbon as a global estimate for terrestrial vegetation (Cramer et al., 2009) to 330 g(H_2_O)/g(C) in the FLUXNET database (Baldocchi and Peñuelas, 2019). This means that considerable amounts of water are necessary for sustaining the basic functioning of plant communities, and particularly of forests which can grow high leaf areas (Asner et al., 2003) leading to high evapotranspiration fluxes. Being sessile organisms, plants rely on the soil stores to provide this water resource. Since the amount of water stored in the soil and accessible to plants is generally lower than the amount of water that could be evaporated considering the local energy balance, plant communities have evolved a range of controls of the water loss to the atmosphere from the scale of the leaf (i.e. stomatal closure; Martin-StPaul et al., 2017) to the canopy (i.e. control of the leaf area index; Eagleson, 1982). More generally, the modification of a large range of physiological processes (Hsiao et al., 1976) along the depletion of the plant water potential illustrates the adaptation of plants to water shortage. Located at both ends of the soil-plant-atmosphere water continuum, the atmospheric demand for water vapor (that relates to vapor pressure deficit, VPD), and the capillary forces retaining water molecules in the soil impose a tension on the water-column ascending the plant xylem. Increases in the atmospheric demand and/or the tension of water in the plant result in progressive stomatal closure (Martin-StPaul et al., 2017) that tends to mitigate the water potential drop-down and prevent xylem embolism.

Characterising the response of the canopy to VPD is rather straightforward (Oren et al., 1999; Novick et al., 2016; Grossiord et al., 2020) since atmospheric VPD can readily be measured, or calculated from measurements of relative humidity and temperature. On the other hand, the role of soil moisture in controlling canopy gas exchanges has remained more difficult to quantify. Soil matric potential (in MPa), which would be the physical measurement most relevant to quantify the soil water availability to the plants, is rarely measured. Instead, soil water content (SWC, in 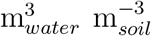) is usually monitored, and then possibly interpreted as water potential through pressure-volume functions (e.g. van Genuchten, 1980). Even then, SWC is usually measured over shallow soil depths for practical reasons (typically down to 20-50 cm, see e.g. Granier et al., 2007; Novick et al., 2016), which in many cases is not suitable for quantifying the total stock of water available to trees. Indeed, trees often grow roots deep in the soil, down to several meters (Fan et al., 2017), which can be essential for provisioning the water needed for transpiration and the overall tree functioning during periods of water shortage (Germon et al., 2020; Christina et al., 2017; McCormick et al., 2021) even if representing a small fraction of the root system mass (Jackson et al., 1996). Specifically in oaks, Lucot and Bruckert (1992) documented the presence of fine roots down to 4 meters in a *Quercus robur* plot with no physical or chemical constraints. Bréda et al.(1995) reported both the presence of fine roots and water extraction down to 2-meters in *Quercus petraea*.

Reflecting this “shallow-soil bias”, models of ecosystem functioning have so far mostly considered soil water to be extractable only in the upper horizons of the soil, defining the so-called “soil water holding capacity” (SWHC, the amount of water, in millimeters, extractable from the soil by plants) at typical depths of one to two meters (e.g. Piedallu et al., 2011; Krinner et al., 2005; Dufrene et al., 2005; Granier et al., 2007). This parameter has been shown very sensitive in the modelling of ecosystem water balance (Granier et al., 1999), carbon balance (Dufrene et al., 2005), tree growth (Guillemot et al., 2017) and survival (Cheaib et al., 2012; Preisler et al., 2019). Challenging the SWHC concept, and in accordance with the recognition of deep water as essential for the functioning of trees, recent works have defined the Total Available Water (TAW) concept, that adds “deep water” extraction by trees to SWHC considered over 1-m (Carriere et al., 2020). However, this deep water resource remains poorly quantified because the trees actual rooting depth and the capacity of deep roots to extract water are not known.

Improving our understanding of soil water provisioning of forests is particularly relevant in the context of climate change, because terrestrial ecosystems are facing an increase in atmospheric evaporative demand (Grossiord et al., 2020) and projections of reduced summer precipitation point to a likely increase in edaphic water stress over the coming decades in most continents and notably Western Central Europe and the Mediterranean zone (Samaniego et al., 2018). How forests react to increased water deficit will depend, to a large extent, on their access to soil water.

Recent summer drought and heat events observed over large parts of Europe (year 2003 in Western Europe, 2010 in Russia, 2018 in Northern central Europe; Bastos et al., 2020) have constituted natural experiments documenting the influence of warmer and drier conditions on ecosystem carbon and water fluxes. In particular, the combined summer 2003 drought and heat caused a seasonal reduction of gross and net carbon uptake (Reichstein et al., 2007; Bastos et al., 2020), as well as of evapotranspiration, in response to soil water stress (Granier et al., 2007).

As the return period of combined drought and heat events decreases with ongoing climate change (Samaniego et al., 2018), it is urgent to improve our understanding of the provisioning of water to forest ecosystems, and to evaluate the sensitivity of modelled projections of ecosystem functioning to the parameterization of soil properties.

Here we used soil water content data measured over a 150-cm soil profile, combined with deep soil water content estimates (i.e. sensors down to −300 cm) to evaluate the role of surface vs. deep water in provisioning tree transpiration needs and modulating canopy gas exchange in a temperate oak forest. We focused on the two contrasted, consecutive years of 2017 (a mild summer) and 2018 (characterized by a hot drought). Our objectives were:

1. to evaluate the response of a temperate oak forest to a hot drought in terms of carbon and water balances,
2. to estimate the Soil Water Holding Capacity (SWHC, in millimeters of water) of the forest,
3. to estimate the contribution of soil horizons to the provision of water for canopy transpiration, contrasting both a dry and milder year,
4. to evaluate how the combinations of SWC and VPD modulate canopy conductance and transpiration,
5. to compare two estimates of SWHC: one provided by soil databases vs. one derived from our measurements of SWC, and quantify the sensitivity of the carbon balance simulated by a process-based model to the choice of one or the other SWHC estimate.

## Materials and Methods

### Study site

The Fontainebleau-Barbeau site (FR-Fon, Delpierre et al., 2016) is located in the Paris area, near the Fontainebleau forest (48.476358 N, 2.780120 E, 103 m a.s.l.). The site has a gentle slope of 2°towards the Seine river which flows 750 m at the South-West of the flux tower. Sessile oak (*Quercus petraea* (Matt.) Liebl.) is the main species, accounting for 79% of the basal area and dominating an understorey mostly occupied by hornbeam (*Carpinus betulus* L., 18% of the basal area). The dominant height is 28 m. The soil is an endostagnic luvisol (IUSS Working Group WRB, 2015) covered by an oligo-mull humus. It developed on a calcareous bedrock located at 5-10 m-depth (Thiry, 2010) and has a discontinuous buhrstone layer located between 60 to 90-cm depth, over which a perched water table develops in winter, usually till late spring. Mean annual air temperature and cumulated precipitation are 11.2°C and 677 mm, respectively (1980–2010, Melun weather station, 12 km from the study site). The leaf area index (LAI) was on average 6.01 over the 2012–2018 period. In spring 2021, we established the root distribution at FR-Fon by counting roots according to diameter class in a 2-m wide and 150-cm deep trench dug 50 meters away from the flux tower. We noticed the presence of fine roots down to the bottom of the trench (−150 cm; Suppl. Mat. S1). In autumn 2021, we extracted one 5-m deep core sample, also in the vicinity of the flux tower. On this unique sample of 10-cm diameter, we observed the presence of 1-mm diameter roots down to a depth of 4 meters (Suppl. Mat. S1), at the top of a dense green marl layer.

### Eddy covariance and micrometeorological measurements

We measured the fluxes of CO_2_ and H_2_O between the forest and the atmosphere with the eddy covariance (EC) method, using both a Li7500 open-path and a Li7200 enclosed-path analyzer (LI-COR Inc., Lincoln, NE, USA), associated with a R3-50 and a HS-50 sonic anemometer (Gill Instruments Ltd., Lymington, UK), respectively. Both EC systems runned concurrently at 37 m a.g.l., i.e. 9 m above the dominant height. EC data were acquired at 20 Hz with two data loggers time-synchronized with a PTP server: a CR3000 (Campbell Scientific Inc., Loughborough, UK) for the Li7500/R3-50 and the Smartflux 2 for the Li7200/HS-50 (LI-COR Inc., Lincoln, NE, USA). Half-hourly fluxes of CO_2_ and H_2_O were calculated using the EddyPro software (v7.0.6, LI-COR Inc., Lincoln, NE, USA). Frequency response corrections were applied to raw fluxes, accounting for high-pass (Moncrieff et al., 2005) and low-pass filtering (Moncrieff et al., 1997). The influence of air density fluctuations on the fluxes was accounted for (Burba et al., 2008). As usually done (Baldocchi, 2008), we did not correct eddy covariance fluxes despite a 18±6% average energy imbalance (calculated as the Energy Balance Ratio, Wilson et al., 2002) at the FR-Fon site. Half-hourly CO_2_ and H_2_O fluxes have been acquired continuously from 2005 by the Li7500/R3-50 device and were considered the reference data series here. Fluxes computed from the Li7200/HS-50 device, running from 2012, were used to gap-fill the reference data through linear regressions established annually. Following this instrumental gap-filling, the calculated Net Ecosystem Productivity (NEP) and evapotranspiration (ETR) fluxes were quality-controlled, gap-filled, and NEP was partitioned into Gross Primary Productivity (GPP) and ecosystem respiration (Reco) according to standard procedures (Reichstein et al., 2005; Papale et al., 2006). We estimated the uncertainty of annual GPP and NEP values as the combination of uncertainties originating from the choice of a u* threshold for filtering the data, and the propagation of random error inherent to EC measurements during the statistical inference by the gap-filling algorithms (Delpierre et al., 2012). ETR is an integrative measurement of the quantity of water evaporated from both wet surfaces (soil and canopy) and plant transpiration. Here we partitioned the eddy-covariance ETR flux into (i) the canopy plus topsoil litter evaporation flux, that do not affect the amount of water held in the organo-mineral horizons of the soil and (ii) the transpiration plus soil evaporation flux (TrSeF), that tap the soil water stock. For doing so, we applied to eddy-covariance ETR data the ratio of TrSeF to ETR calculated hourly by the CASTANEA model (see e.g. Nelson et al., 2018). Air temperature and relative humidity used to calculate the vapor pressure deficit (VPD) were measured at 37 m a.g.l with a HMP155A thermohygrometer (Vaisala Corp., Helsinki, Finland) protected from radiation with a 43502H aspiration radiation shield (R. M. Young, Traverse City, MI, USA). Precipitation was measured with a ARG100 raingauge (EML, North Shields, UK raingauge) located at 36 m a.g.l.

### Soil water content

#### Data acquisition

Soil water content (SWC, in 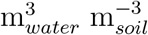) was measured from 2012 with five EnviroSCAN probes (Sentek Sensor Technologies, Stepney, Australia) located in the vicinity of the tower and connected to a CR1000 datalogger. Each probe had a 10-cm vertical distribution of 15 sensors from −5 cm to −145 cm, i.e. down to 150 cm depth. From 2016, a sixth probe reached the depth of −300 cm with the following vertical sensor distribution: −15, −55, −85, −135, −175, −215, −255, and −295 cm. The CR1000 datalogger achieved automatic data acquisition with half-hourly means calculation. In order to assess the spatial representativity of SWC measurement collected by the 6 probes, 24 access tubes of 160 cm long were installed in the vicinity of the tower in 2012 over a 3800-m^2^ area. We measured SWC in those tubes on a weekly basis, with a DIVINER probe (Sentek Sensor Technologies, Stepney, Australia), measuring SWC in the soil profile at multiple depths (at 10 cm intervals), comparable to the profiles obtained with the EnviroSCAN probes. The EnviroSCAN and DIVINER measurements are based on high-frequency capacitance. Since no soil-specific calibration of the probes was achieved on the Fontainebleau-Barbeau site, SWC were calculated from the scaled frequency of the probes using the factory calibration equation. We tested the sensitivity of the calibration equation, following the work of Provenzano et al. (2016), and found minor influence on the SWC dynamics, in accordance with observations by the probe manufacturer (Sentek Pty Ltd, 2001).

#### Estimation of the Soil Water Holding Capacity

We estimated the integrated SWHC down to 300 cm using the following equation:

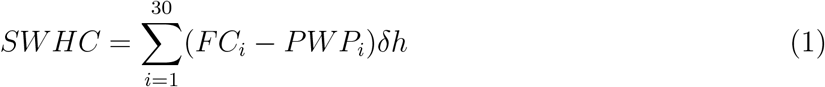

with the index i identifying soil layers of δh = 10 cm depth down to 300 cm. We identified FC (Field Capacity, i.e. the maximum moisture of unsaturated soil, in 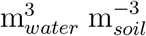) for each soil layer, by looking at periods of stable, relaxed SWC following soil saturation occurring typically occurring in early spring. For each soil layer, we identified the permanent wilting point (PWP, in 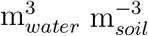) as the lowest SWC value for that soil layer, typically reached during dry summer periods. As SWC drops to PWP, root water uptake, and the associated daily oscillations of SWC, tend to zero in the considered soil layer. We illustrate the basic procedure to extract FC and PWP for the layer between 120 and 130 cm (index i=13) in Fig.2, assuming the PWP is reached in the very dry year 2018. For each layer in the 0-160 cm zone, FC and PWP have then been deduced from the SWC closest sensors. The highest vertical resolution (equal to 10 cm) and representativeness of the spatial variability (20 probes) is obtained for indexes i= 1 to 16 (i.e. 10 to 160 cm depth). Less information was available for the deeper layers (160-300 cm). In that case, a unique automatic probe measuring SWC at 180, 220, 260, and 300 cm has been used to investigate soil layers i= 17 to 30. As a consequence, and because we suspected those deep soil layers to remain above PWP in year 2018 (see below), we used FC and PWP determined over i=14 to 16 as representative of soil layers i= 17 to 30 (Suppl. Mat. 4).

**Fig. 1:**
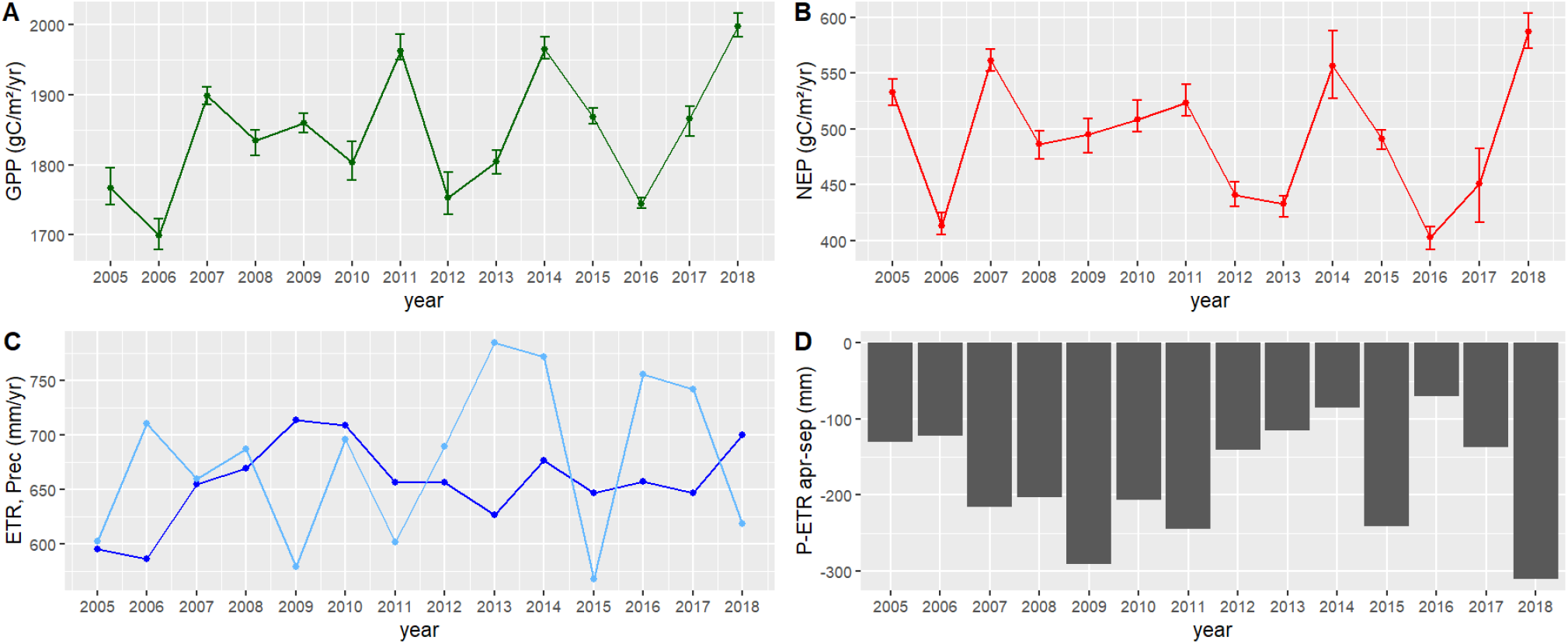
Annual carbon and water fluxes at FR-Fon for the period 2005-2018. (A) displays the annual gross primary productivity (GPP), (B) the net ecosystem productivity (NEP), (C) the annual precipitation (Prec, light blue) and evapotranspiration (ETR, dark blue) and (D) a climatic water balance, namely the difference between Apr-Sep precipitation and evapotranspiration. In (A) and (B), error bars display the 5th and 95th percentile of the GPP and NEP uncertainty distributions (see text for details).

**Fig. 2:**
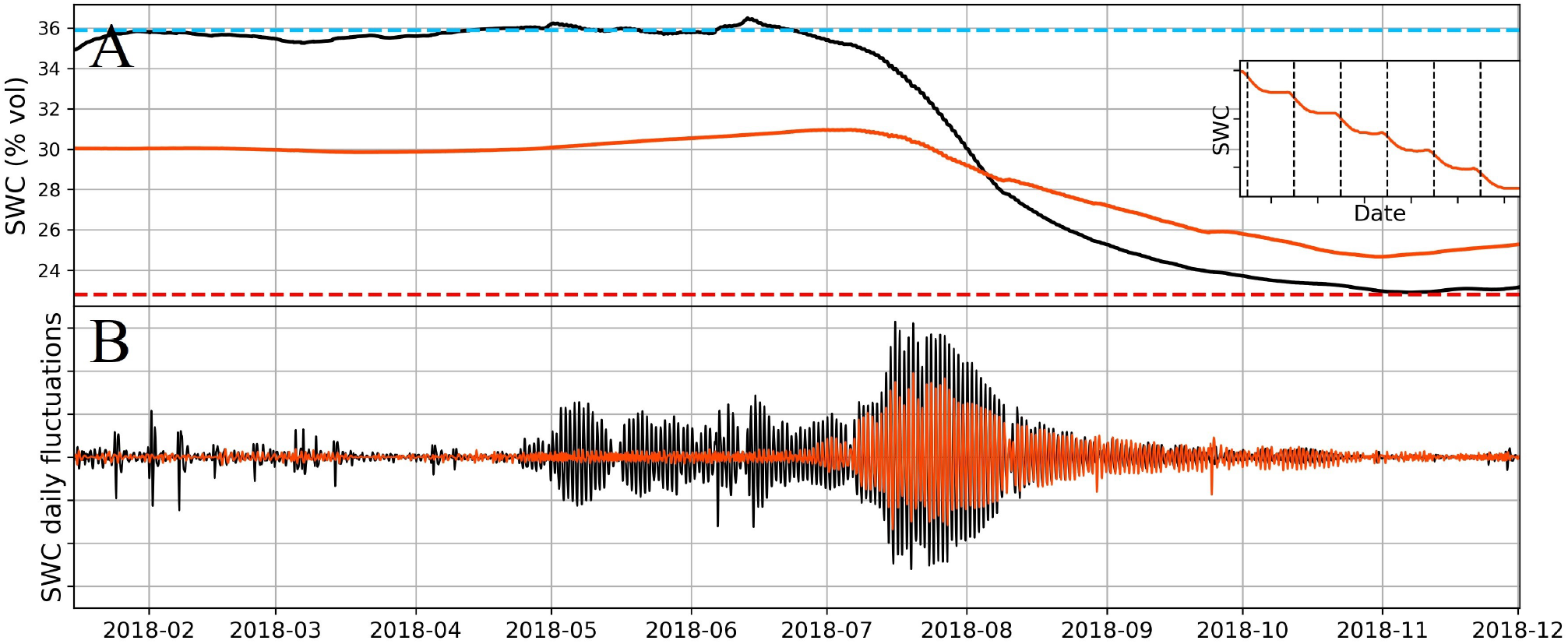
SWC time series of the layer 120-130 cm (black color) and 290-300 cm (orange color) during the 2018 dry year. (A) displays the measured SWC profile calculated as the mean of five 160-cm length EnviroSCAN probes (for layer 120-130 cm); one 300-cm length EnviroSCAN probe (for layer 290-300 cm). The skyblue- and red-dashed lines are the FC and PWP values for layer 120-130 cm, respectively. (B) displays results from a passband filter (fourth-order Butterworth filter) applied to the SWC profiles and centered around daily fluctuations. The inset in (A) is a zoom on SWC profile at 290-300 cm between 2018-08-02 and 2018-08-08, with vertical dashed lines indicating midday.

#### Estimation of SWHC uncertainty

We assumed the SWHC uncertainty to result from the reading of PWP and FC on the SWC time series and the spatial (among-probes) variability. The reading uncertainty was estimated at σ_*r*_= 1 mm for PWP and FC respectively. The uncertainty σ_*s*,*i*_ attributed to spatial variability can be accurately estimated for each 10-cm layer i thanks to the DIVINER probe exploring a 3800-m^2^ area over 160 cm (see above). For deeper (170-300 cm) layers, we hypothesized σ_*s*,*i*_ was equal to the average value in the upper zone. As a consequence, we calculated the the overall SWHC uncertainty as:

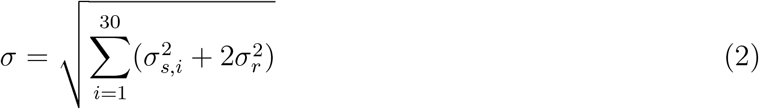

where i index denotes the layer number ranging from 1 (layer 0-10 cm) to N=30 (layer 290-300 cm).

### Calculation of Root Water Uptake

The amount of water uptaken from the soil by trees, the so-called RWU flux (Root Water Uptake, in mm day^−1^), can be calculated from SWC time series (Hupet et al., 2002). The major interest of using SWC data to infer RWU is that it does not require any prior knowledge of the root distribution. In that context, different methods exist to extract RWU, from numerical model inversion (Zuo and Zhang, 2002) to the use of daily fluctuations in SWC (Li et al., 2002). Indeed, RWU manifests as fluctuations in soil moisture in the rhizosphere, influenced by evapotranspiration which occurs preferentially over daytime. As a consequence, RWU can be assessed using linear regression and differentiation of daytime vs. nighttime change in SWC. In a methods comparison (Guderle and Hildebrandt, 2015), this approach using daily SWC oscillations proved to be the best regarding a combination of criteria: the quality of estimations and lesser induced error due to moisture sensor accuracy (the calibration error). Consequently, we focused on this method, and calculated RWU at our site from SWC measurements with the python package *rootwater* (Jackisch and Malicke, 2020). This algorithm analyses daily fluctuations in SWC and takes into account possible hydraulic redistribution, a mechanism by which water is moved via the root system from moister to drier soil layers (Caldwell et al., 1998; Ishikawa and Bledsoe, 2000; Neumann and Cardon, 2012). We applied the *rootwater* algorithm on days when SWC profiles exhibited the specific transpiration-induced shape of RWU (i.e. daytime decline of SWC). We therefore rejected days displaying strong drainage along the soil profile, which could blur the effect of RWU. For selected days, a reference daily moisture profile was built, based on the SWC values at sunset/sunrise, and linear interpolations. The reference reflects “perfect” RWU over daytime and diffusion over night time (Jackisch et al., 2020). The theoretical step shape profile was then compared to the observed profile thanks to an evaluation of the Nash-Sutcliffe efficiency (NSE, Nash and Sutcliffe, 1970).

### Response of canopy conductance to VPD and SWHC

In order to evaluate how the combinations of SWC and VPD modulated canopy conductance (G_*can*_) and evaporation, we first calculated G_*can*_ through inverting the Penman-Monteith equation, as in Novick et al. (2016). We then fitted G_*can*_ as a function of VPD with the following equation (Novick et al., 2016):

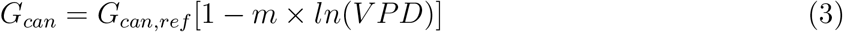

where G_*can*_ is the canopy conductance for water vapor (mmol m^−2^ s^−1^), G_*can*,*ref*_ is its value at reference VPD of 1 kPa, and *m* (ln(kPa)^−1^) is the slope of the relation. We fitted this equation for different classes of soil humidity, computed as Relative Extractable Water (REW, defined as REW = SWC_0–300*cm*_/SWHC_0–300*cm*_ where SWC_0–300*cm*_ is the actual stock of water extractable by trees, in millimeters, over a 300-cm soil depth for a given day and SWHC_0–300*cm*_ is the maximum water content extractable by trees, in millimeters, over a 300-cm soil depth).

### CASTANEA model

One of our objectives was to determine the sensitivity of ecosystem models to parameterizations of SWHC. To this aim, we used the CASTANEA model (Dufrene et al., 2005) to simulate CO_2_(namely GPP) and H_2_O (namely ETR) fluxes at FR-Fon, under different climate conditions. CASTANEA is a process-based ecosystem model that simulates the forest carbon and water fluxes (including transpiration, canopy, litter and soil evaporation) at an half-hourly time step. First, we calibrated some of the model most sensitive parameters (Supplementary notes SN1) in order to simulate GPP and ETR accurately over periods with no water stress over the period of 2006-2019. Then, we performed prospective simulations under a changing climate, considering three climate models under two RCP scenarios (RCP4.5 and 8.5, see Jourdan et al., 2021 for details on the climate models). With those simulations, we quantified the sensitivity of simulated GPP to the SWHC parameter. SWHC is a forcing parameter to the model, and has been identified as very sensitive, with 10% variation of SWHC yielding 13-24% variations of annual NEP, 5-8% variations of annual GPP and 3-4% variations of annual ETR in simulated fluxes (Dufrene et al., 2005).

### Soil database for parameterizing SWHC

We parameterized CASTANEA with two different SWHC estimates at the FR-Fon forest. We used our own estimate of SWHC, based on SWC measurements (see Results). Beside this, and similar to what is usually done for parameterizing ecosystem models (e.g. Cheaib et al., 2012), we used a SWHC estimate obtained from a soil database (Badeau et al., 2010). Badeau et al. built a 1-km^2^ resolution database of SWHC over France, from a soil database of soil texture and rooting depth (Jamagne et al., 1995). Each 1×1 km grid point encompasses different soil types, and the database gives for each grid point the average value of SWHC with respect to the area of each soil type over the 1×1 km grid.

## Results

### Annual fluxes and climatic water balance

The growing season (April-September) climatic water balance (P-ETR) was largely negative on all years (−179 mm on average) and reached its lowest value in 2018 (−310 mm; Fig.1d). In spite of this, year 2018 was the year of highest GPP (1997 gC m^−2^ y^−1^, Fig.1a) and the year of highest NEP on record (587 gC m^−2^ y^−1^, Fig.1b), as compared to averages of 1844 gC m^−2^ y^−1^ and 491 gC m^−2^ y^−1^ respectively over 2005-2018. Annual evapotranspiration averaged 657 mm y^−1^ over 2005-2018, and reached its third highest value in 2018 (700 mm y^−1^, Fig.1c). Accordingly, when considering seasonal patterns, GPP, NEP and ETR appeared not influenced by the climatic water deficit evolving over the 2018 growing season (Suppl. Mat. S3 and S5).

### Estimation of SWHC

In order to estimate SWHC, we focused on year 2018, which was the driest on record (based on the climatic water balance data, Fig. 1d; see also the comparison of 2017-2018 soil water content data, Suppl. Mat. S5). In that year, we observed seasonal variations of soil water content down to 300 cm, which was the location of the deepest SWC probe. In the 290-300 cm layer, SWC fluctuated from 0.31 to 0.25 m^3^/m^3^ during year 2018, reaching its maximum in July and its minimum in November (Fig.2a). The seasonal dynamics at 290-300 cm was clear, although less ample than at a shallower layer (e.g. 120-130 cm, Fig.2a) where it fluctuated from 0.36 (end of June) down to 0.23 m^3^/m^3^(November, Fig.2a). Interestingly, we observed daily fluctuations of the SWC signals at all measurements depths, down to 290-300 cm (Fig.2b). The daily fluctuations were neat from May at 120-130 cm depth and their amplitude increased in July, when they also became visible in the 290-300 cm layer (Fig.2b). As for the upper layers (not shown), the SWC signal had a clear shape in the 290-300 cm layer, with a decrease during the day and a stall during the night, which was evocative of the day/night cycle of tree transpiration, and root water uptake (Fig.2a inset).

We estimated the quantity of water extractable in each soil layer from SWC measurements (Suppl. Mat. S4). Field capacity (FC, the maximum moisture of unsaturated soil) was considered to be reached when SWC reached stable, high values (e.g. skyblue dashed line on Fig.2a), after checking the disappearance of a seasonal perched water table from piezometer data (not shown). We supposed that one soil layer had reached the permanent wilting point (PWP) when SWC reached a stable, low value (e.g. red dashed line on Fig.2a). We only had one probe for characterizing the water availability deeper than 160 cm (measuring at depths 170-180, 210-220, 250-260 and 290-300 cm, cf. Material and Methods), as opposed to 20 probes for shallower layers (0-160 cm). Below 160 cm, it was also difficult to identify clear FC and PWP values (e.g. for the 290-300 cm layer on Fig.2a). Hence we extrapolated values of FC and PWP obtained for the layers 140-160 cm (Suppl. Mat. S4) to deeper layers, which is coherent with the homogeneous nature of the soil (sandy loam) over 120-300 cm. We calculated SWHC to reach 390 mm over 0-300 cm, with an estimated uncertainty of 17 mm, combining the reading uncertainty of FC and PWP, and the variability observed among the 20 measurements probes.

### Seasonal and interannual patterns of root water uptake

The root water uptake (RWU) flux calculated from the *rootwater* algorithm applied over 0-300 cm compared generally well with the transpiration fluxes derived from ET measurements (Fig.3). Both the seasonal shapes, and the actual flux values were similar between those independent methods. The *rootwater* algorithm tended to generate lower flux values as compared to the ET-derived transpiration (average of 2.22 vs. 2.34 mm/day, respectively, in 2017 and 2.34 vs. 2.56 mm/day in 2018) though the extrema were similar (minima of 0 and maxima of 6-7 mm per day). From Figure 3 we inferred that water extraction could occur deeper than 300 cm since RWU (calculated over 0-300 cm) was generally lower than the ET-derived transpiration flux and appeared noticeably underestimated in year 2018 on DoY 240-260, when extraction from deep horizons was particularly strong (see below).

**Fig. 3:**
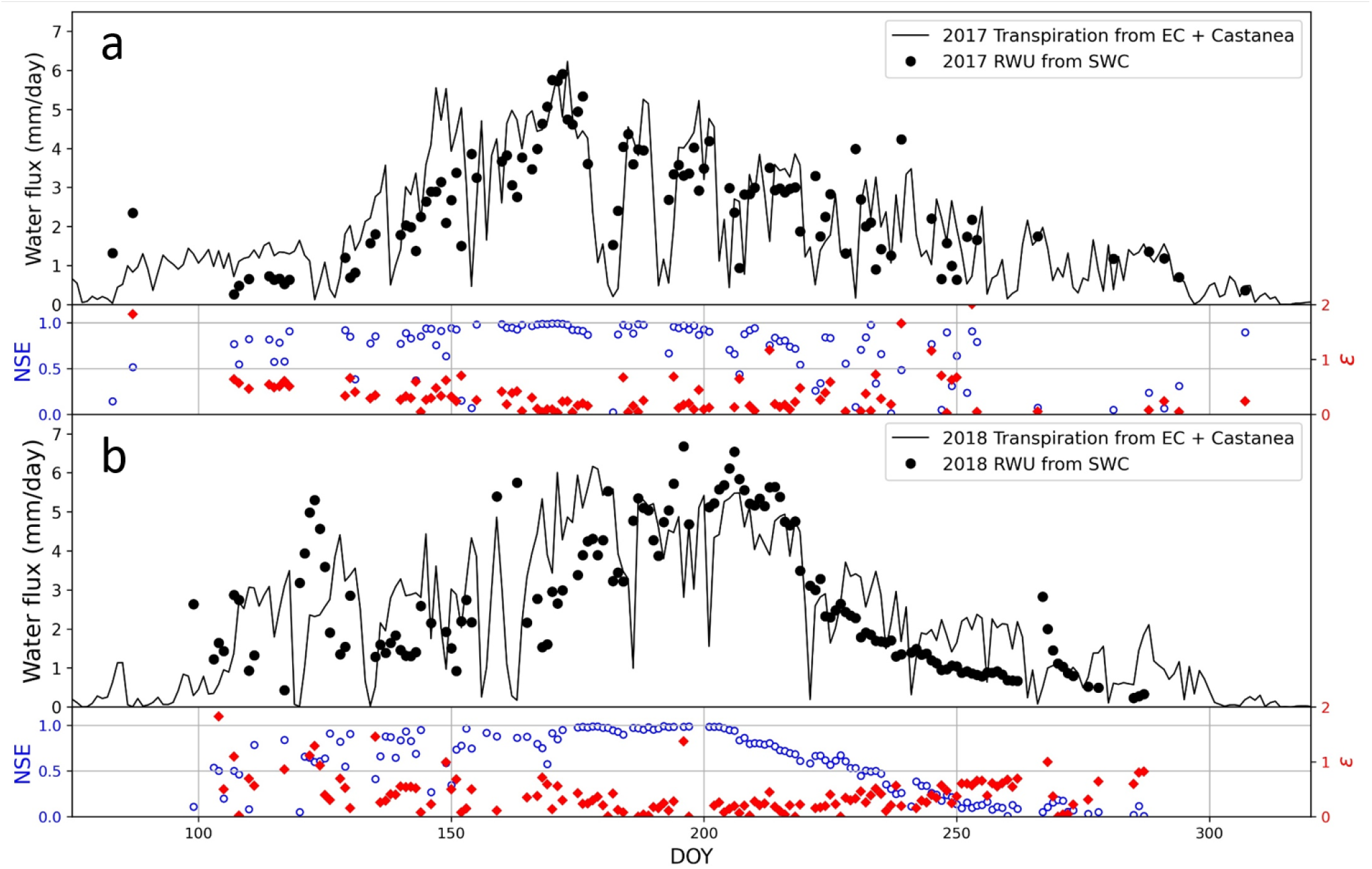
Daily water fluxes in mm per day over the year in 2017 (a) and 2018 (b). Black dots illustrate the root water uptake (RWU) calculated from the soil water content (SWC) integrated over 300 cm, assuming no hydraulic redistribution. The black-solid line corresponds to tree transpiration (T) resulting from the daily average evapotranspiration fluxes calculated from the flux tower (EC), from which T was deduced thanks to the CASTANEA ecosystem model (see text). Red markers represent the absolute relative error between the water fluxes estimated from SWC and EC which is obtained as *ε* = abs(RWU-T)/T. Blue dots represent the daily NSE values of the *root water* algorithm applied over 0-300 cm.

The seasonal patterns of RWU in individual soil layers are displayed on Fig.4. The contribution of deep layers increased as superficial layers dried up. The plasticity of RWU was fast. For instance, the contribution of the 0-150 cm layers decreased from 72% to 55% over 12 days (DoY 165 to 176 in 2017, Fig.4a); from 69% to 51% over 7 days (DoY 214 to 220 in 2017, Fig.4a); and from 64% to 43% over 21 days (DoY 198 to 217 in 2018, Fig.4b). More generally, the role of deep water extraction increased over the growing season. Before DoY 160, the contribution to RWU of layers deeper than 150 cm was less than 20% in both years. This contribution went up to more than 40% and almost 60% on DoY 220, for 2017 and 2018 respectively.

**Fig. 4:**
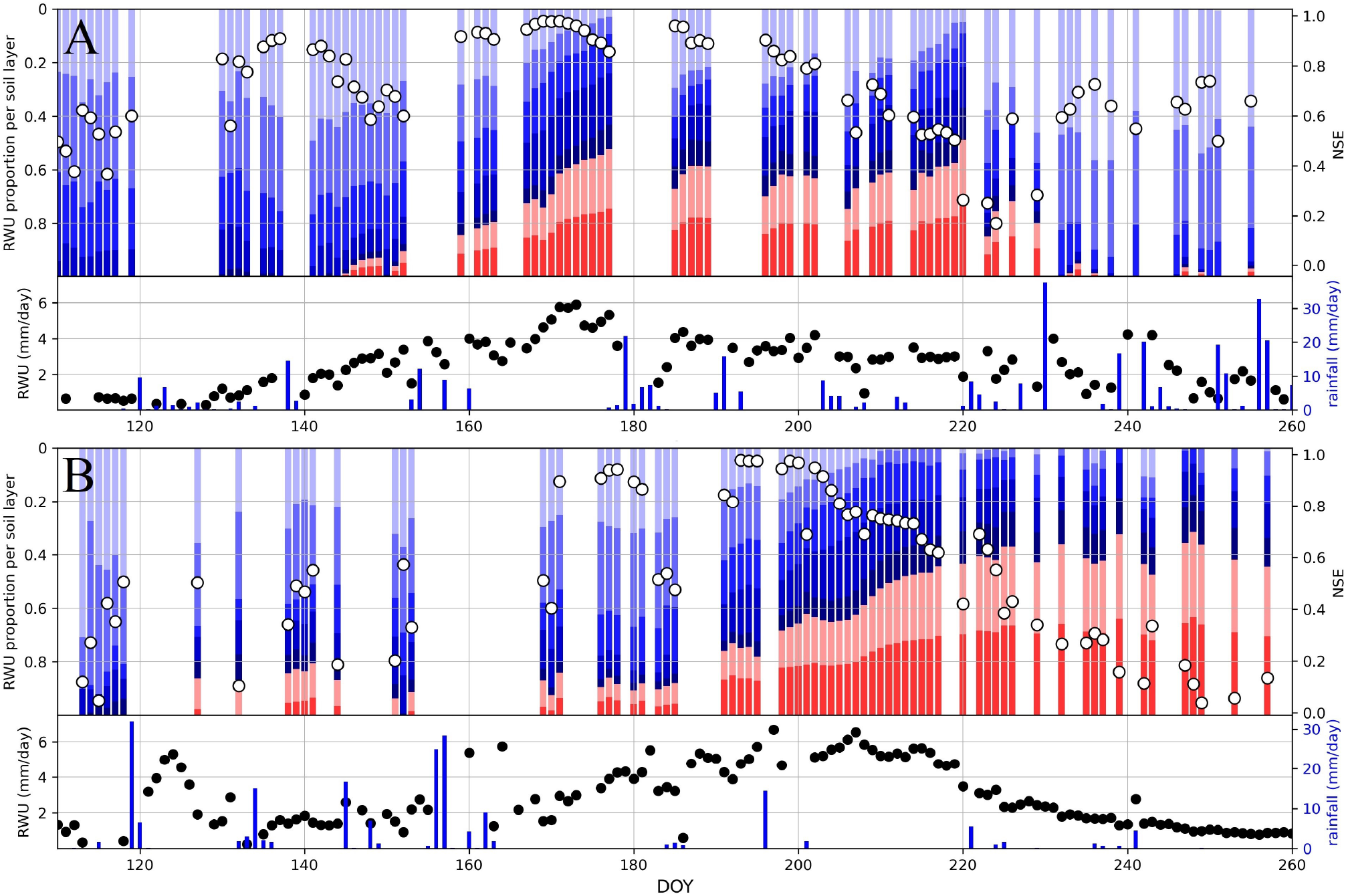
Root water uptake (RWU) dynamics in 2017 (a) and 2018 (a). The RWU in mm day^−1^ was estimated using the *rootwater* algorithm on the integrated soil water content considered over 0-300 cm depth. The proportion per horizon was obtained using the information on separated layers, assuming hydraulic redistribution if observed on the SWC profile. The colors corresponded to different layers, from lightest to darkest blue: 0-30 cm, 30-60 cm, 60-90 cm, 90-120 cm and 120-150 cm depth. Layers 150-220 cm and 220-300 cm are in light red and dark red respectively. Five EnviroSCAN probes were used to calculate RWU over 0-150 cm, one probe was used over 150-300 cm (see text). White dots (right axis) represent NSE values of the *rootwater* algorithm. Blanks in the RWU time series mark episodes of high precipitation (typically above 5 mm per day) hindering the estimation of RWU by the algorithm.

RWU seasonal patterns differed for years 2017 and 2018, owing to their meteorological characteristics. For instance, over the period of DoY 160-180, we observed deeper water extraction in 2017 as compared to 2018, in relation with drier soil conditions (Suppl. Mat. S5). In the same logic, the drier period of DoY 220-260 in 2018 showed a deeper water extraction as compared to 2017.

### Soil influences canopy conductance despite high SWHC

The contribution of deeper layers to RWU increased as soil dryness developed over the growing season. This revealed the plasticity of water uptake to maintain high levels of evapotranspiration. Yet this plasticity was limited by the total amount of water in the soil, as shown in Fig.5. On panels (a) and (b), half-hourly latent heat fluxes were plotted against the Relative Extractable Water (REW) defined as REW = SWC/SWHC_0–300*cm*_, over the period DoY 165-255. REW decreased from 0.6 to 0.2 in 2017 (Fig.5a) and from 1.0 (i.e. field capacity) to 0.1 in 2018 (Fig.5b). The large REW range explored over 2018 made the correlation between latent heat fluxes and REW clearly visible (Fig.5b). Latent heat fluxes showed no trend along REW for high REW values (typically over 0.6) and decreased as REW decreased.

**Fig. 5:**
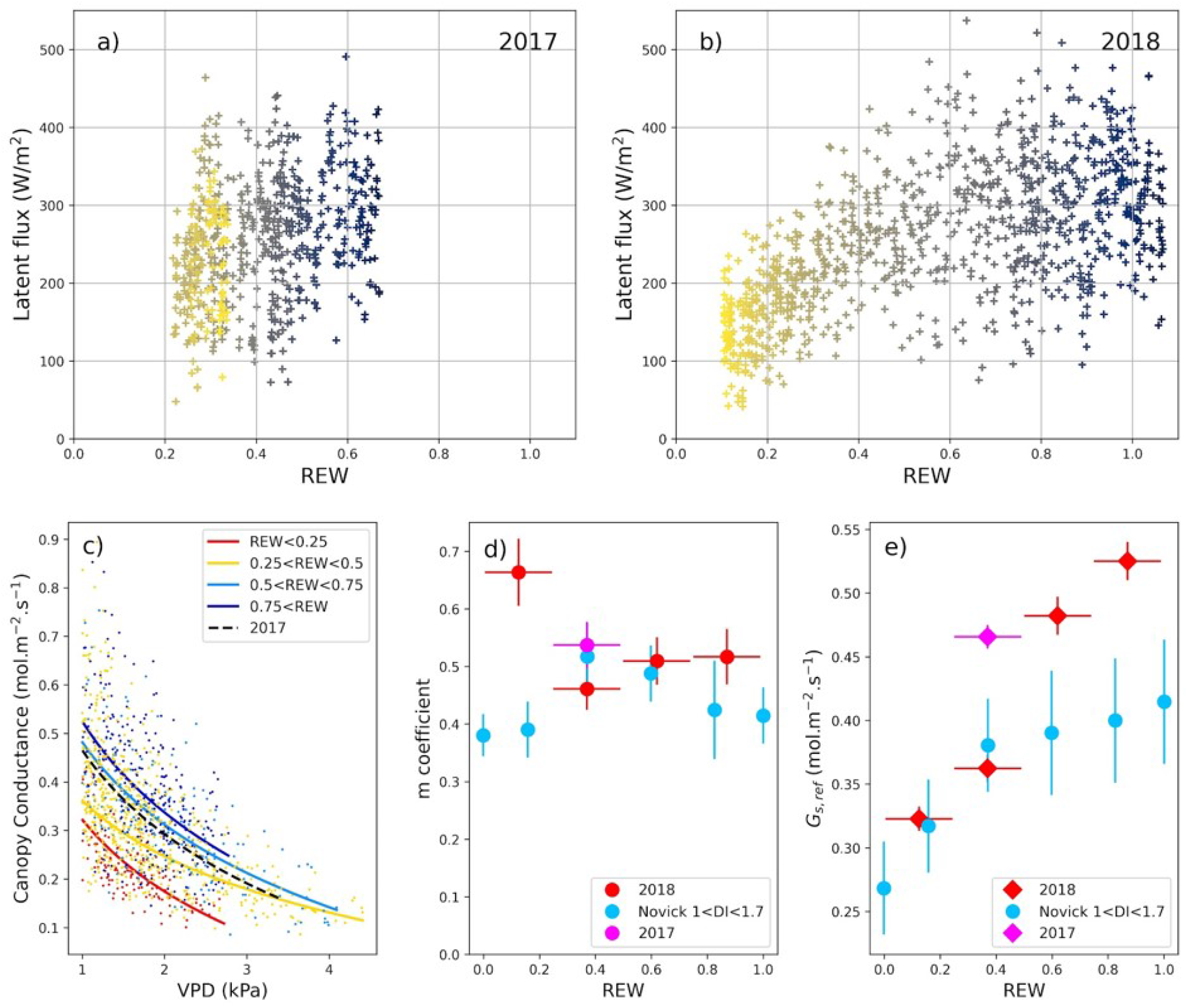
Effect of SWC and VPD on canopy conductance and evapotranspiration. The data plotted were selected in 165<DoY<255, between 10 am and 6 pm, with VPD≥1 kPa and latent heat flux≥150 W/m^2^. In (a) and (b), the point colors refer to DoY, with the darkest points for DoY 165 and the yellowest points for DoY 255. In (c), the lines represent the fit of equation 3 for different intervals of REW, with the colored lines being for 2018 while the black dashed line is for 2017 with 0.25<REW<0.5. Panels d) and e) display the values of *m* and G_*can*,*red*_ calculated from our data as well as those calculated by Novick et al. (2016) (their Figure 2 e and f) over an ensemble of Ameriflux sites. We represented here average values obtained at sites comparable to Fontainebleau-Barbeau according to values of the Dryness Index (the ratio of annual PET to annual P, with values from 1.0 to 1.7). For the sake of comparison, SWC data reported by Novick et al. (2016) (over 0-30 cm) were converted to REW by dividing the actual SWC value by the maximum SWC value reported on their figure.

Figure 5c further illustrated the dual control exerted by REW and VPD on canopy conductance. Canopy conductance decreased with both VPD and REW (Fig.5c). At a reference VPD of 1 kPa, canopy conductance (G_*can*_=G_*can*,*ref*_) decreased from 0.52 to 0.32 mol m^−2^ s^−1^ (Fig.5c) as REW decreased from 0.87 to 0.12 (Fig.5e, showing that the values are comparable to those reported by Novick et al., 2016 for mesic sites), while *m*, the slope of the relation between, G_*can*_ and VPD, was in the range [0.46; 0.66], apparently increasing as REW decreased (Fig.5d, also showing data in line with Novick et al., 2016), noting that the comparison of the *m* slopes along REW may be uncertain because *m* at low soil water content was estimated from a fit established over a narrow VPD range (red curve on Fig.5c).

### Sensitivity of an ecosystem model to SWHC parameterization

The SWHC value of 390 mm determined from SWC measurements was 1.9 times higher than the value of 207 mm estimated from the 1-km French soil database for the corresponding grid point (Badeau et al., 2010), and in the high end of the SWHC distribution of soils appearing in the database over France (Fig.6a). The CASTANEA model fairly simulated the seasonality of GPP observed at FR-Fon when parameterized with SWHC= 390 mm (Fig.6b), as well as the interannual variability of GPP (Suppl. Mat. SN1). On the contrary, parameterizing CASTANEA with SWHC= 207 mm, as obtained from the soil database (Badeau et al., 2010) leaded to higher soil water stress and reduced GPP in summer, as compared to GPP derived from the flux tower measurements (Fig.6b). Over the period 2006-2019, annual GPP simulated with SWHC= 390 mm were 365 gC m^2^ y^−1^ higher than with SWHC= 207 mm. The GPP anomaly caused by the under-estimation of SWHC in the model parameterization remained rather stable in future climate conditions under RCP4.5 scenario, but nearly doubled to 700 gC m^2^ y^−1^ for two out of three climate models under RCP8.5 (Fig.6c).

**Fig. 6:**
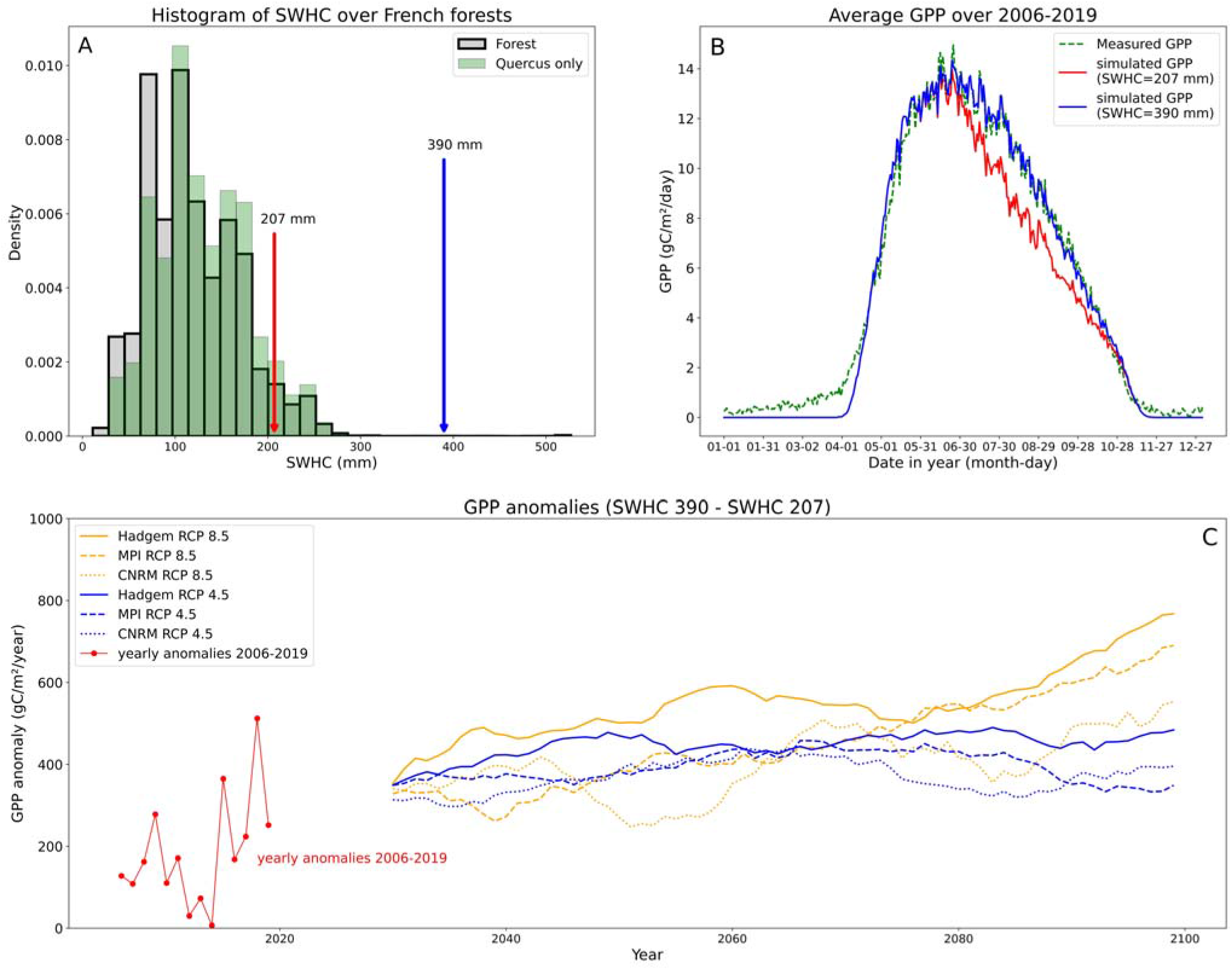
Sensitivity of the CASTANEA model simulations to the parameterization of Soil Water Holding Capacity (SWHC). (a) Distribution of the SWHC values in the French Soil Database, with all 1-km^2^ pixels occupied by forests (grey bars, n= 68 373 points) and forest pixels mostly occupied by deciduous oaks (n= 35 352 points). Red arrow point to the SWHC estimate obtained for FR-Fon grid point in the Badeau et al. (2010) database, Blue arrow point to our estimate of SWHC, obtained from the analysis of SWC measurements; (b) Comparison of GPP calculated from the FR-Fon flux tower measurements with CASTANEA simulations for two values of SWHC (390 mm and 207 mm), averaged over the 2006-2019 period; (c) Projections of simulated difference in GPP between CASTANEA model runs using SWHC= 390 mm and SWHC= 207 mm over the 2020-2100 period (lines displayed are the 20 years rolling averages). Three climatic models were considered (Hadgem, MPI and CNRM) with two CO_2_ emission scenarios for each (RCP 4.5 and RCP 8.5). The red points at the beginning of the time series illustrate the average anomaly calculated over 2006-2019.

## Discussion

### Influence of Drought 2018 on carbon and water fluxes at FR-Fon

The growing season of 2018 was amongst the driest encountered at FR-Fon over the past 40 years (ranked sixth over the period 1980-2018 as regards the difference of P-PET, not shown). Yet the annual productivity remained high, with the highest annual NEP and highest annual GPP from the start of flux measurements in 2005 (Fig.1a,b). Lower than average GPP values over July-October were compensated by higher than average GPP over April-July (Suppl. Mat. S3b), probably owing to higher than average temperatures and radiation (Suppl. Mat. S2a,b). The higher than average spring photosynthesis was not caused by an early budburst, which occurred on DoY 105 for oaks (i.e. equal to the average 2006-2018 date, Soudani et al., 2021). The July-October 2018 negative anomaly of GPP was moderate, probably in relation to the high SWHC. Indeed, a lower SWHC would have resulted in lower GPP (Fig.6b). Higher than average NEP were encountered during most of the growing season, and notably during July-October, when GPP was lower than average. During this period, Reco was strongly reduced (Suppl. Mat. S3c), possibly as consequence of summer drought. This was not in line with the higher sensitivity of GPP than of Reco to soil water stress, as evidenced during the 2003 hot drought (Granier et al., 2007). A similar report of 2018 higher than average NEP and GPP owing to enhanced spring carbon uptake, followed by close-to-average summer values has been reported in a Czech floodplain deciduous forest (Kowalska et al., 2020). More generally, the analysis of an ensemble of DGVM simulations and the FLUX-COM products (i.e. fluxes simulated by machine-learning techniques) over central Europe showed a seasonal (summer) influence of the 2018 heat and drought on carbon fluxes, but little to no influence on carbon fluxes at the annual scale (Bastos et al., 2020), nor on stem growth (Salomén et al., 2022). This contrasted with reports of negative annual carbon uptake anomalies evidenced in Northern Europe (Lindroth et al., 2020).

### Accuracy of the SWHC estimate

We estimated SWHC to reach 390 ±17 mm at FR-Fon. This value was the 99.93^*th*^
 percentile of the distribution of SWHC values estimated with pedotransfer functions integrated down to 1-meter for French forests or oak stands (i.e. the same percentile for both distributions, Fig.6a). It was also very high when compared to estimates published at other sites, at which roots have been evidenced to go deeper than 1 meter (e.g. 165 mm for a 200 cm root exploration depth in an oak stand in Bréda et al.,1995, maximum of 186 mm over 16 European forest sites with soils reaching depths down to 160 cm in Granier et al., 2007, see also De La Motte et al., 2020). However, SWHC values similar to, or even higher than our estimate of 390 mm have been published for temperate forests (e.g. 338 mm for the Hesse Beech forest in North-Eastern France, De La Motte et al., 2020, or 450 mm for a mixed Beech-Oak forest near Louvain-la-Neuve in Belgium, De Wergifosse et al., 2020). The value of 390 mm for SWHC was 1.9 times higher than the database estimate of 207 mm for the FR-Fon grid point (Badeau et al., 2010). It is noticeable that the latter estimate was obtained from the integration of pedo-transfer functions over a soil depth of 112 cm, far lower than the actual root exploration depth at FR-Fon (Suppl. Mat. S1). If considering a soil depth of 112 cm, we would estimate a SWHC of 191 mm based on our SWC measurements (integration of SWHC data per soil layers displayed in Suppl. Mat. S4). From this, we conclude (1) that our estimate of SWHC is conservative, meaning that deriving FC and PWP from SWC data (Suppl. Mat. S4) yielded similar (actually, slightly lower) values of SWHC than obtained from pedo-transfer functions (Badeau et al., 2010); (2) that what makes the SWHC of FR-Fon forest so high as compared to Badeau et al. (2010) grid point estimate is the difference of root exploration depth. Contrasting with the 112 cm-depth estimate of Badeau et al. (2010), our analysis of SWC data suggest that roots can actually extract water at least down to −300 cm at FR-Fon (Fig.4), which is supported by the observation of roots down to −300 and even −400 cm on one soil core taken at the site (Suppl. Mat S1).

Actually, we consider the 390-mm value as a lower-bound estimate for SWHC at FR-Fon. Indeed, we estimated this value on the basis of SWC measurements recorded during the dry 2018 summer. While we are confident that our estimates of water content at field capacity were reliable, we may have underestimated the values of the wilting point (Suppl. Mat. S4), particularly for the soil layers below 100-cm depth that may have remained above the wilting point during this dry episode (note the decreasing trend of SWHC with depth below 100-cm in (Suppl. Mat. S4a). A body of evidence further supports this hypothesis: (i) our calculation of Root Water Uptake (RWU) is slightly lower than the ETR flux measured by eddy covariance (Fig.3). Yet, the latter is possibly underestimated in relation with the 18% lack of energy balance closure at the FR-Fon site. Had we increased ETR by 18% to close the energy balance, our current estimate of RWU would have more neatly lagged behing ETR. (ii) We observe that even at low REW (REW<0.25, red curve on Fig.5c), the canopy conductance remains relatively high. For instance Granier and Bréda (1996) (their Fig.1), report in a sessile oak stand values of canopy conductance (G_*can*,*max*_)of 1 cm s^−1^(= 0.41 mol m^−2^ s^−1^) under high water supply at VPD= 1 kPa, comparable to our estimates of 0.50 mol m^−2^ s^−1^ at FR-Fon (Fig.5c and e). Their value of G_*can*_ decreases to 60% of G_*can*,*max*_ for REW= 0.2 (Fig.2 in Granier and Bréda, 1996). In our case, we estimated that G_*can*_ decreased to 63% of its maximum value (0.33 as compared to 0.52, Fig.5e) for REW= 0.1, pointing to a possible underestimation of REW that would stem from the underestimation of SWHC.

### Temporal pattern of RWU extraction along the soil profile

The *rootwater* algorithm (Jackisch and Mälicke, 2020) fits a model to the daily variations of SWC in a given soil layer to calculate root water uptake. In the original publication, the authors consider a model fit of NSE>0.5 as indicative of most reliable RWU estimates (Jackisch et al., 2020). Our RWU calculations did not always meet this criterion, with some values of NSE<0.5 notably in early summer (before DoY 120 in 2017, or DoY 170 in 2018) and late summer (after DoY 220 in 2018) (Fig.3), meaning that processes other than root water uptake (i.e. water percolation, capillary rise) may intervene. Yet we are confident that our calculations of RWU, and their distribution among soil layers are representative of the actual uptake of water by roots, not only because RWU integrated over 0-300 cm matched the canopy transpiration flux (Fig.3) but also because the algorithm achieved model fits with NSE>0.5 at times when the deepest layers (150-300 cm) contributed a large amount of RWU (e.g. DoY 160-220 in 2017 and up to DoY 220 in 2018, Fig.4). Our calculations of root water uptake (RWU) showed that deep soil layers (i.e. below 150 cm) provisioned some water for canopy transpiration early during the leafy season, from DoY 150 (end of May) in 2017 and DoY 130 (mid-May) in 2018 (Fig.4). This contribution increased substantially, up to achieving 50% of the water uptaken for transpiration over DoY 180-220 in 2017, and even 60% over DoY 210-260 in 2018 (Fig.4). A similar pattern of increasing contributions of deep soil layers to water provision for tree transpiration has already been reported (e.g. Bréda et al., 1995 in an oak site, Bréda et al., 2002 in an ash forest). It is however not systematic and, for instance, a regular, not increasing, and minimal contribution of soil layers below 170 cm depth was reported in a Beech forest along a mild summer in Germany (Jackisch et al., 2020).

Interestingly, Bréda et al., 1995 reported, as we do, variations of SWC in soil layers located below the known root depth (reaching −200 cm in their study). We notice here that significant variations of SWC were observed at a depth of 3 meters (Fig.2b), i.e. 150 cm below the root depth established from root counting profiles made at our site (Suppl. Mat. S1). Yet, the root profile did not “close” at −150 cm at FR-Fon, and we found punctual evidence (from one soil core of 5.5-m length) that roots can actually grow down to −300 cm, and even −400 cm at this site. These observations were coherent with rooting depth of −200 cm, which are frequently reported in *Quercus petraea* (Bréda et al., 1995; Lebourgeois and Jabiol, 2002) and we notice that that fine roots down to −400 cm have already been reported in *Quercus robur* (Lucot and Bruckert, 1992).

### Underestimating SWHC yields substantial errors in the simulation of the forest productivity

Predicting the effects of current and future water stress on the functioning and distribution of forests is one of the key issues in ecological research. A variety of mechanistic models can be used to this aim, with some hypothesizing the forest vulnerability to rely foremost on its carbon balance (e.g. Dufrene et al., 2005), some others simulating the integrity of the plant hydraulics (Cochard et al., 2021), and some integrating both (Davi and Cailleret, 2017; Naudts et al., 2015). Whatever the model type, it requires an accurate estimate of the amount of soil water that can be used by trees. Several papers have pointed recently the role of “deep” water reserves as key to sustain ecosystem functioning (McCormick et al., 2021; Fan et al., 2017; Carriere et al., 2020). We showed that using a SWHC estimate of 390 mm, as opposed to 207 mm as predicted by a reference soil database, impacts strongly the simulation of GPP in the CASTANEA model (Fig.6c). CASTANEA is a “carbon-centric”, still sensitive to the soil water balance through a modulation of the stomatal conductance vs. assimilation relation (Dufrene et al., 2005). This model is not able to simulate the degree of xylem embolism, expected to increase as climate warms and dries (Cochard et al., 2021). Running a hydraulic-enabled model such as SUREAU (Cochard et al., 2021) or MAESPA (Duursma and Medlyn, 2012) at FR-Fon would quantify to what extent the deep-component of SWHC (i.e. the extra 390-207= 183 mm of soil water reserve evidenced in this paper) helps mitigating the expected increase of embolism, and probability of death of trees in a warmer and drier climate.

## Conclusions and perspectives

We evidenced that the FR-Fon oak forest can tap at least 390 mm of water from the soil. This is a large amount for a temperate forest, as compared to published estimates of SWHC based on measurements (Bréeda et al., 1995; De La Motte et al., 2020) or the modelling (Granier et al., 2007) of the soil water balance. This value is 1.9 times higher than the estimate of SWHC (= 207 mm) retrieved for the FR-Fon grid point in the French consensus soil database (Badeau et al., 2010), which is used in modelling studies predicting the future of French forests (Cheaib et al., 2012). Yet, several papers pointed recently the role of “deep” water in provisioning water to sustain ecosystem functioning, particularly during drought periods as the one encountered at FR-Fon in 2018. The monitoring of soil water has generally received little attention at flux measurements sites, with measurements conducted at most sites over shallow depths (typically down to 30-50 cm, Novick et al., 2016). For instance, the ICOS-network protocol states “the minimum depth of the […] SWC profiles is set to 1 m. If limited by the presence of the bedrock or a water-impermeable layer, the profiles can be less deep”(Op De Beeck et al., 2018). Evidence is accumulating (our results and e.g. McCormick et al., 2021; Carriere et al., 2020) that water from deep soil layers can be extracted by trees in many situations, even in the presence of a “bedrock or water-impermeable layer”, and that the SWHC has so far been generally largely underestimated in forests (Fig.6a), at least locally. Hence we stress here that the setup proposed by Op De Beeck et al. (2018) should be considered a minimum. SWC measurements conducted over too shallow depths will overlook the provisioning of water for canopy evapotranspiration by deeper layers. The contribution of these deep layers will probably increase as drought become more frequent and intense with climate change. While installing probes deep in the soil can be difficult, or even not practicable for some rocky sites, indirect methods such as electrical resistivity tomography (e.g. Carriere et al., 2020) are promising for evaluating the rooting depth and getting improved, higher estimates of SWHC. Beyond the documentation of SWHC at particular sites, we need more accurate maps of SWHC in order to improve the simulation of the functioning and vulnerability of forests under climate change.

## Supporting information

Supplementary Material to "Contribution of deep soil layers to the transpiration of a temperate deciduous forest"

## Acknowledgements

We thank Augusto Zanella and Cécile Quantin for their contributions in the description of soil cores, Kamel Soudani and Eric Dufrene for insightful discussions and their critical readings of the manuscript, Claude Doussan for discussing with us questions relative to the role of capillarity, and students Annaёlle Desgrippes and Jocelyn Ragu for their contribution in the acquisition of root distribution data. IC acknowledges funding by the ANR under the “Investissements d’avenir” programme with the reference ANR-16-CONV-0003 (CLAND).

## Authors contributions

ND, JM and DB designed the research; DB, AM and GV collected the data; JM, ND and DB analysed the field data; CF did the model simulations; CF and ND analyzed the model simulations; JM, ND and DB wrote a first working paper; ND wrote the final version of the manuscript with significant inputs from MJ, IC, CF and SB.

