## Supplementary Material to "Contribution of deep soil layers to the transpiration of a temperate deciduous forest" for "Contribution of deep soil layers to the transpiration of a temperate deciduous forest: quantification and implications for the modelling of productivity"

Maysonnave, Delpierre et al.

**Figure S1. Distribution of roots according to diameter class at the FR-Fon forest.**

**Figure S2. Meteorological time series, FR-Fon 2005-2018.**

**Figure S3. Flux time series, FR-Fon 2005-2018.**

**Figure S4. Estimates of Soil Water Holding Capacity for soil layers down to 160 cm at the FR-Fon forest.**

**Figure S5. Time series of soil water content over the whole soil profile (0-300 cm), and daily precipitation in 2017 and 2018 at FR-Fon.**

**Supplementary Note S1. Calibration of the CASTANEA model.**

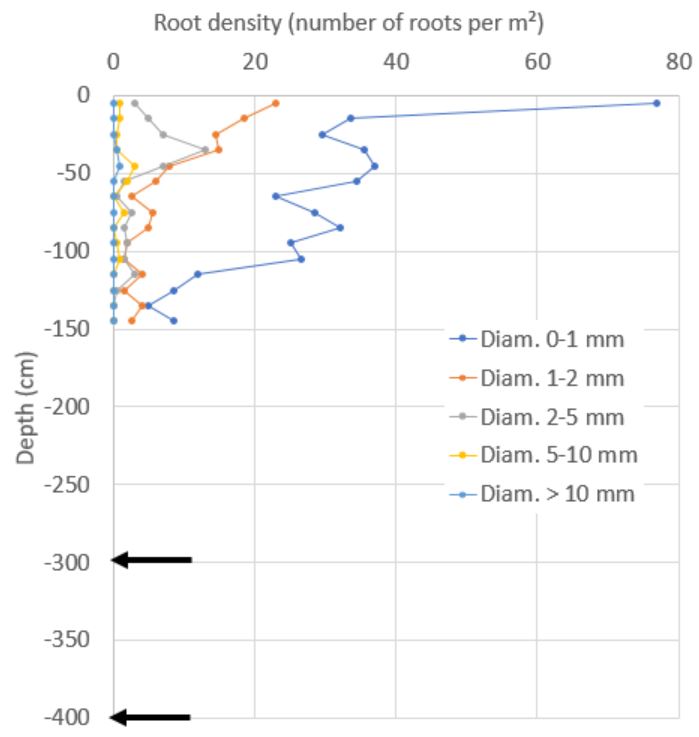

**Figure S1. Distribution of roots according to diameter class at the FR-Fon forest**. The data were acquired along a 200-cm wide, 150-cm deep trench, by counting the number of roots according to diameter class in 10x10 cm² areas. Data were acquired on May 4-7, 2021. Black arrows point to the observed presence of 1-mm diameter roots at -300 and -400 cm, along one 5-m depth soil core of 10-cm diameter, sampled at FR-Fon in October 2021.

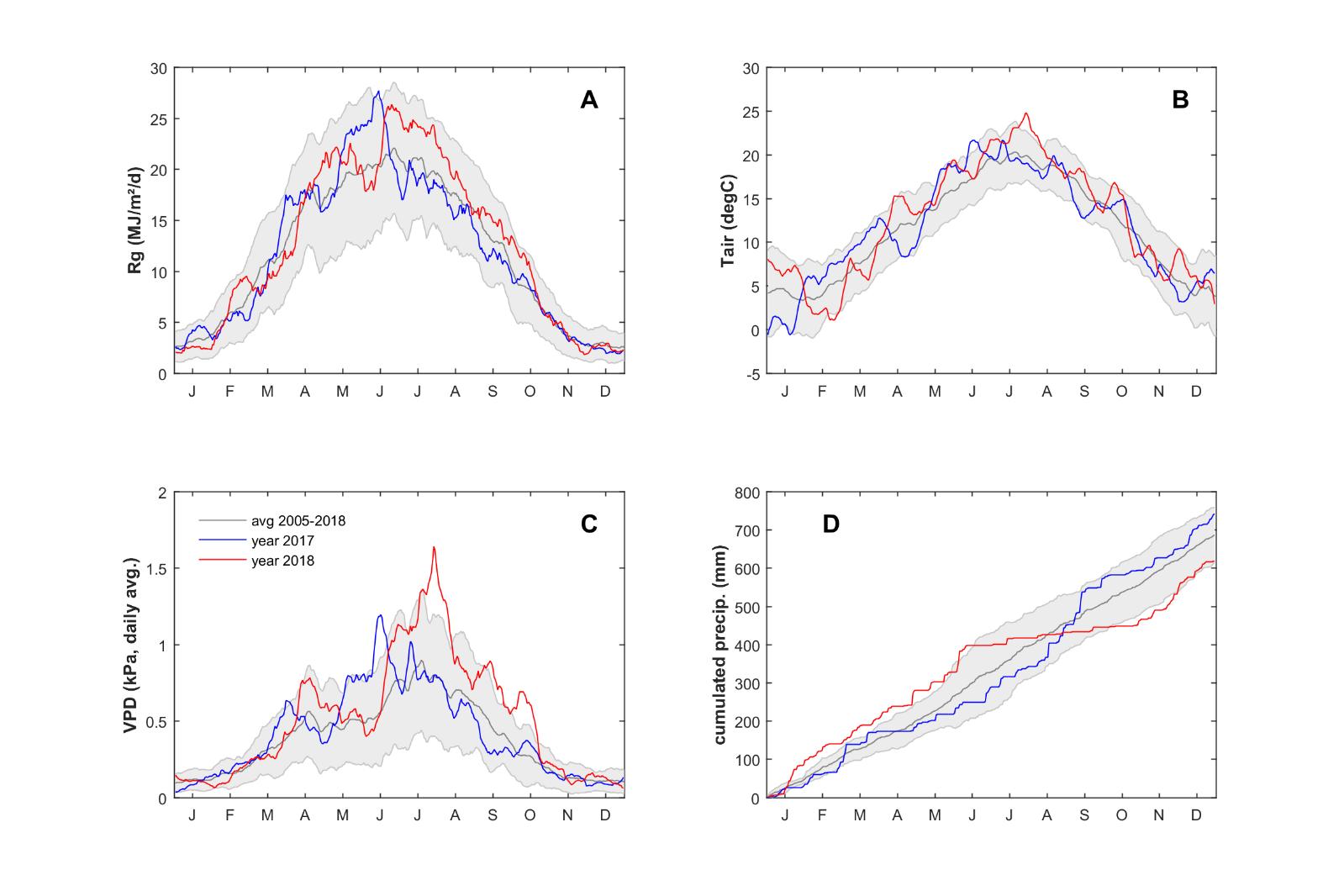

**Figure S2. Meteorological time series, FR-Fon 2005-2018.** Time series of global radiation (A), air temperature (B), VPD (C), and cumulated precipitation. Shown are 2005-2018 average ± one SD (grey), year 2017 (blue) and year 2018 (red). For clarity, the 2005-2018 average patterns are smoothed with a rolling mean of 10 days; years 2017 and 2018 are smoothed with a rolling mean of 15 days.

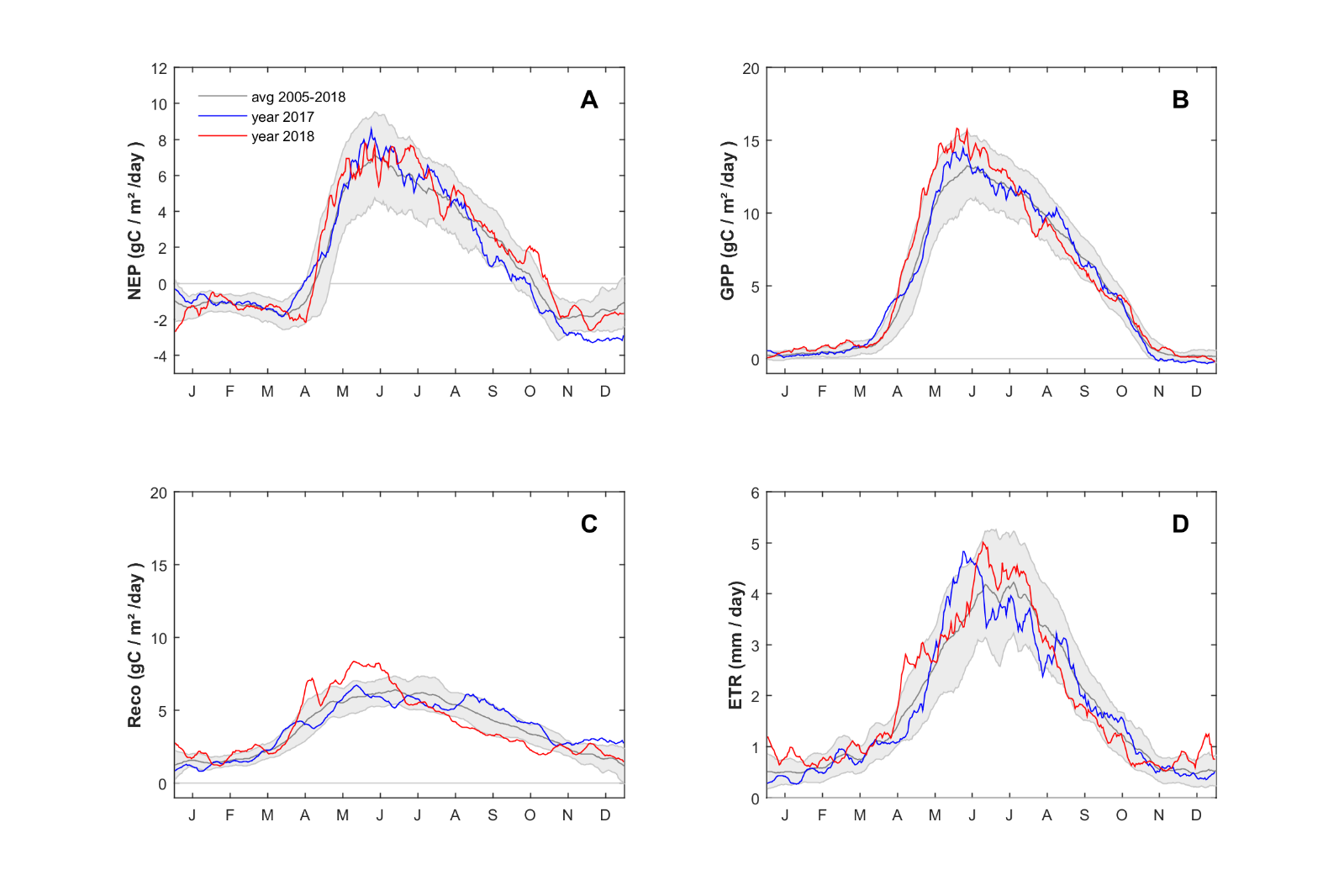

**Figure S3. Flux time series, FR-Fon 2005-2018.** Time series of NEP (A), GPP (B), Reco (C) and ETR (D). Shown are 2005-2018 average ± one SD (grey), year 2017 (blue) and year 2018 (red). For clarity, the 2005-2018 average patterns are smoothed with a rolling mean of 10 days; years 2017 and 2018 are smoothed with a rolling mean of 15 days.

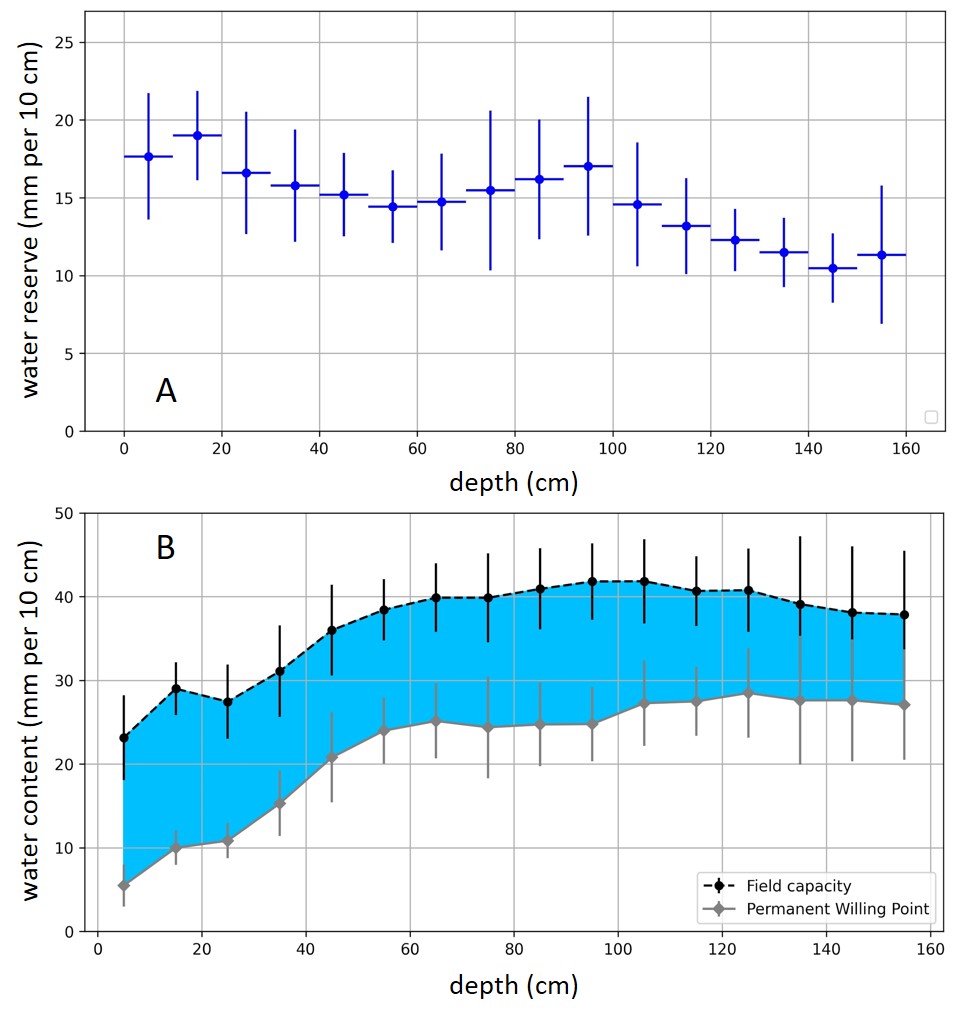

**Figure S4. Estimates of Soil Water Holding Capacity for soil layers down to 160 cm at the FR-Fon forest.** (A) reports the values of SWHC in mm of water per 10 cm soil. (B) illustrates the values of field capacity and wilting point read from the SWC sensors and used to calculate SWHC in (A).

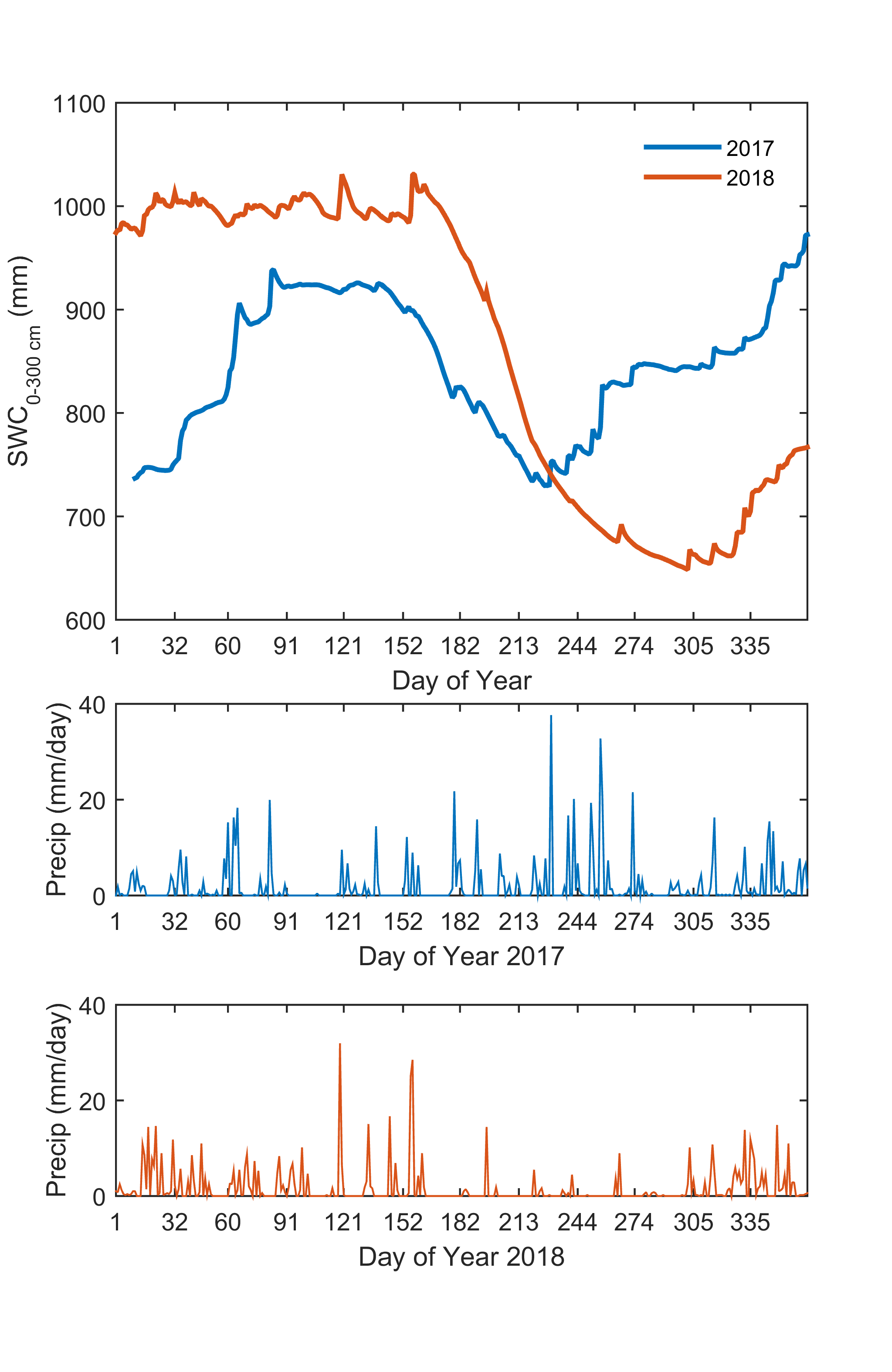

**Figure S5. Time series of soil water content over the whole soil profile (0-300 cm), and daily precipitation in 2017 and 2018 at FR-Fon.**

**Supplementary Note S1. Calibration of the CASTANEA model**

We calibrated several parameters of the CASTANEA model (Dufrêne et al. 2005) on GPP and ETR flux data measured at FR-Fon during the period of 2006-2019.

We focused on three parameters, known to be particularly sensitive (see Dufrêne et al., 2005 for a sensitivity analysis of CASTANEA):

*θ*, the curvature of the quantum response of the electron transport rate (dimensionless)

*g1_max_*, the maximum value of the slope of the Ball et al. model of stomatal conductance (dimensionless)

*α_Na_*, the proportional dependency between the leaf maximal carboxylation rate *VC_max_* and leaf nitrogen content (µmol CO_2_ gN^−1^ s^−1^)

Each parameter has been varied between a minimum and a maximum value (see Table SN1.1) and daily simulated GPP and ETR were recorded between 2006 and 2019. A *score* (resp. GPP_score_ and ETR_score_) has been calculated for each variable (resp. GPP and ETR) as the sum of squares of the differences between measured and simulated variables. The final *total score* was a weighted sum of both scores : *total score* = GPP_score_ + *a* * ETR_score_ where *a* has been set to 4 to take into account the difference of magnitude between GPP and ETR and the different number of selected days (see below). The choice of factor *a* did not have a great importance for the final parameters choice, given the relatively flat form of the *score* functions (see Fig. SN1.1).

Since we wanted to quantify the influence different Soil Water Holding Capacity (SWHC) values on the modelled response of GPP to water stress, we calibrated the above-mentioned three parameters only during days without water stress. Since the simulated GPP is also pretty sensitive to phenological parameters which were not the focus of this paper, we calculated the GPP score for parameter calibration from 42 days after budburst (i.e. at a time when the simulated canopy has reached full development in terms of leaf area and photosynthetic capacity) up to DoY 320 (mid-november). This represented about 2800 days for the 2006-2019 period. In order to avoid known errors in the measurements of ETR with the eddy covariance method during rainy days, we computed ETR scores for parameter calibration only for days without rain, during the leafy season (between days 135 and 250). This represented about 900 days for the 2006-2019 period.

The total score function (for GPP and ETR) over 2006-2019 has been evaluated for about 20 000 (θ, g1_max_ , α_Na_) triplets according to the specified ranges and steps (see Table SN1.1).

The best triplet minimizing the distance between simulated and observed GPP and ETR for the 2006-2019 period is given in Table SN1.1.

**Table SN1.1 Original values for the three calibrated parameters, explored range (minimum and maximum values and step) and final calibrated values for *Quercus petraea* at FR-Fon.**

|  | Original Value | Min tested value | Max tested value | Step | Calibrated Value |
| --- | --- | --- | --- | --- | --- |
| θ | 0.7 | 0.5 | 1 | 0.05 | 0.9 |
| g1_max_ | 9.5 | 7 | 15 | 0.25 | 11 |
| α_Na_ | 30.8 | 15 | 40 | 0.5 | 20 |

Fig SN1.2 shows the result of the calibration for the GPP and ETR simulation over the 2006-2019 period.

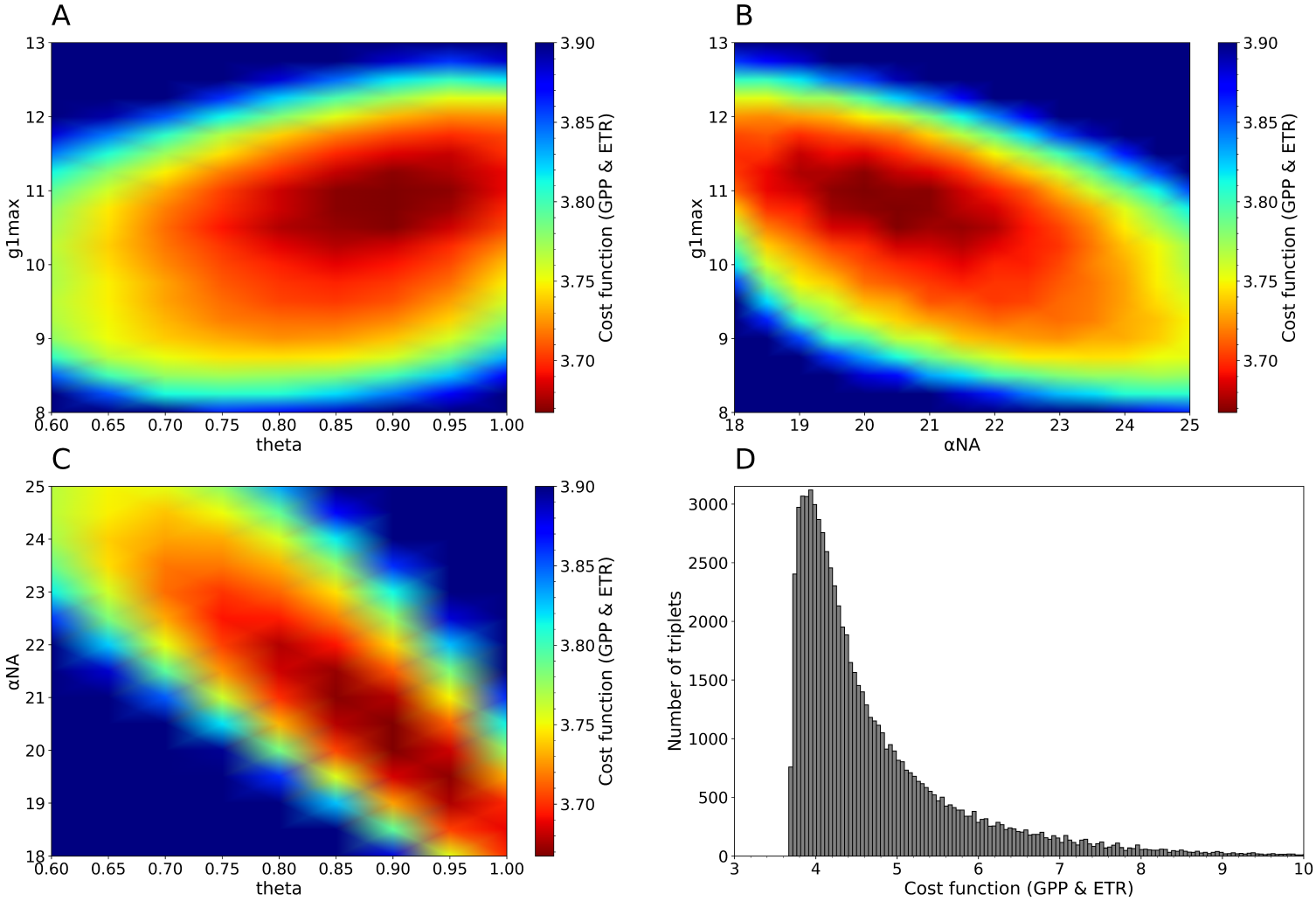

**Figure SN1.1**. A, B, C: graphs of the *total score* function. Colors represent the minimum of the *total score* function for the values of the two represented parameters, regardless of the third parameter. Dark red represents the region of the best values. D: histogram of all the values taken by the *total score* function for the 20 000 triplets.

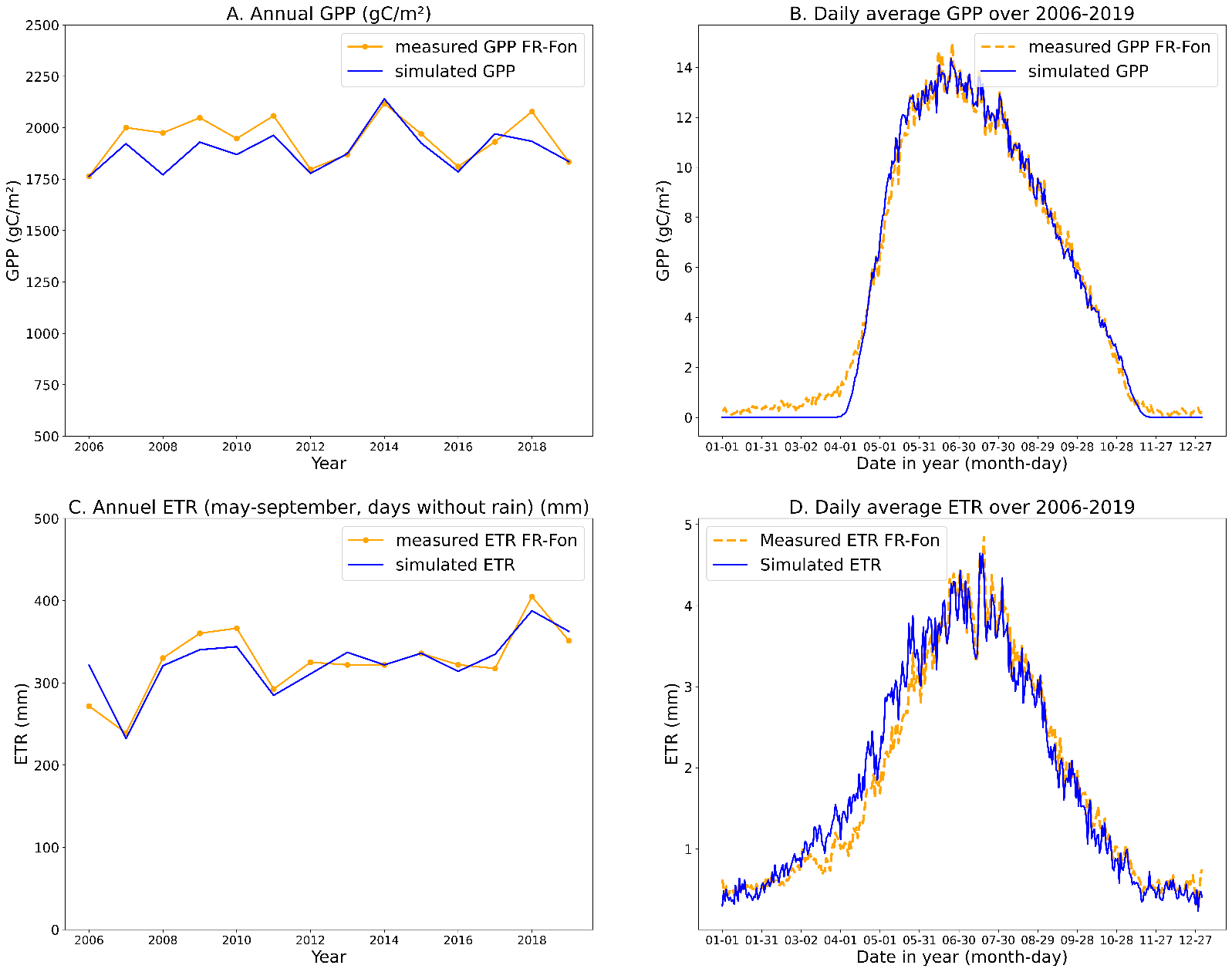

**Figure SN1.2**. Result of the CASTANEA calibration over the FR-Fon data (2006-2019) for GPP (A, B) and ETR (C,D). The calibration was performed for days without water stress over the leafy season. The results presented here include all days of the year (except for C, May-September, days without rain only).
